# An ultrapotent RBD-targeted biparatopic nanobody neutralizes broad SARS-CoV-2 variants

**DOI:** 10.1101/2021.12.25.474052

**Authors:** Xiaojing Chi, Xinhui Zhang, Shengnan Pan, Yanying Yu, Tianli Lin, Huarui Duan, Xiuying Liu, Wenfang Chen, Xuehua Yang, Qiang Ding, Jianwei Wang, Wei Yang

## Abstract

The wide transmission and host adaptation of SARS-CoV-2 have led to the rapid accumulation of mutations, posing significant challenges to the effectiveness of vaccines and therapeutic antibodies. Although several neutralizing antibodies were authorized for emergency clinical use, convalescent patients derived natural antibodies are vulnerable to SARS-CoV-2 Spike mutation. Here, we describe the screen of a panel of SARS-CoV-2 receptor-binding domain (RBD) targeted nanobodies (Nbs) from a synthetic library and the design of a biparatopic Nb, named Nb1-Nb2, with tight affinity and super wide neutralization breadth against multiple SARS-CoV-2 variants of concern. Deep-mutational scanning experiments identify the potential binding epitopes of the Nbs on the RBD and demonstrate that biparatopic Nb1-Nb2 has a strong escape resistant feature against more than 60 tested RBD amino acid substitutions. Using pseudovirion-based and trans-complementation SARS-CoV-2 tools, we determine that the Nb1-Nb2 broadly neutralizes multiple SARS-CoV-2 variants, including Alpha (B.1.1.7), Beta (B.1.351), Gamma (P.1), Delta (B.1.617.2), Lambda (C.37), Kappa (B.1.617.1) and Mu (B.1.621). Furthermore, a heavy chain antibody is constructed by fusing the human IgG1 Fc to Nb1-Nb2 (designated as Nb1-Nb2-Fc) to improve its neutralization potency, yield, stability and potential half-life extension. For the new Omicron variant (B.1.1.529) that harbors unprecedented multiple RBD mutations, Nb1-Nb2-Fc keeps a firm affinity (KD < 1.0×10^−12^ M) and strong neutralizing activity (IC_50_ = 0.0017 nM). Together, we developed a tetravalent biparatopic human heavy chain antibody with ultrapotent and broad-spectrum SARS-CoV-2 neutralization activity which highlights the potential clinical applications.

## Introduction

The current emerging severe acute respiratory syndrome coronavirus 2 (SARS-CoV-2) causes global pandemic and the coronavirus disease (COVID-19) related deaths had exceeded 5.3 million in December 2021^1,2^. The continuing circulation and evolution of SARS-CoV-2 in human and susceptible animals pose a huge challenge to public health and social interaction^3,4^. Clinical manifestations of COVID-19 in the general population range from asymptomatic infection, fever, dry cough, loss of taste or smell to severe pneumonia, multi-organ failure, and death^1,5^. Progress has been made in SARS-CoV-2 small molecule direct antiviral agents by targeting viral RNA-dependent RNA polymerase and main protease (3CL pro)^6-8^. Nevertheless, potent and specific antivirals targeting diverse mechanisms for either prevention or therapy are still urgently needed in the context of a pandemic.

Prophylactic vaccines against SARS-CoV-2 were developed from the multiple technology routes^9-11^, with a major purpose to elicit neutralizing antibodies. However, vaccines are unable to protect individuals with low immunity, autoimmune diseases, and low vaccination willingness. The emergency use authorization (EUA) has been issued for clinical utility of neutralizing antibodies to treat certain COVID-19 patients^12,13^. As the critical function for binding to the host receptor ACE2 and cell entry^14^, the receptor-binding domain (RBD) on SARS-CoV-2 Spike protein is the most preferred antigen target for neutralizing antibody-based countermeasures^15-17^. The antigenic landscape of the SARS-CoV-2 RBD can be divided into seven binding communities, including the receptor binding motif (RBM), the outer face of the RBD, and the inner face of the RBD^18^. Neutralizing antibodies binding to RBM provide the most potent activity, while neutralizing antibodies associated with the outer face of the RBD demonstrate excellent neutralization breadth^18^.

The SARS-CoV-2 is constantly evolving and has accumulated many mutations across its genome, especially within the Spike gene^19^. Distinct variants of concern (VOC) or variants of interest (VOI), such as Alpha (B.1.1.7), Beta (B.1.351), Gamma (P.1), Delta (B.1.617.2) and Omicron (B.1.1.529), are associated with enhancement of virus transmission and jeopardize neutralizing antibody activities through potential diminished or loss of binding^20,21^. The desired neutralizing antibodies require a difficult balance between neutralizing potency and broad-spectrum. This is why the majority of clinical monoclonal antibodies adopt antibody pairs that recognize two or more distinct Spike epitopes, known as “cocktails” strategy. For all this, any single monoclonal antibody has to face the risk of viral escape.

A VHH antibody, also known as nanobody (Nb), is the antigen binding fragment from camelid or shark heavy chain antibody, which is the smallest antibody fragment with antigen affinity^22,23^. Nb alone is about 12-15 KDa, and composed of four conserved framework regions (FRs) and three hypervariable complementarity-determining regions (CDRs). Nb has unique biological and physical features, including low manufacturing cost, prominent stability, adjustable half-life, alternative routes of administration, and prone to synthesizing the homo/hetero multimers from diverse functional Nb building blocks^24^. Evidence suggests that Nbs can exhibit super-strong activity and a broad binding spectrum, through combining different Nbs into a new polyvalent molecule^25^. Therefore, Nbs are becoming a powerful weapon against viral diseases.

In this study, we obtain several SARS-CoV-2 RBD targeting Nbs with either high-affinity or broad neutralization spectrum using a previously developed synthetic nanobody discovery platform^10^. We identified a biparatopic Nb as the best in class broadly neutralizing antibody, which can potently neutralize more than 60 SARS-CoV-2 Spike pseudotyped viruses bearing single point, combination, and deletion mutations, as well as multiple VOC and VOI, including the new super mutant Omicron variant. Collectively, our study has characterized a single antibody, rather than a cocktail of antibodies, with ultra-broad RBD coverage which significantly reduces its risk of viral escape and provides an alternative for optimizing COVID-19 prophylactic and therapeutic antivirals.

## Results

### Selection and design of neutralizing Nbs with potent activity and affinity

The unceasing accumulation of mutations in the SARS-CoV-2 Spike causes the loss of efficacy for some established neutralizing antibodies^26^. Iterative discovery and identification of neutralizing antibodies against emerging variants will provide a solid stockpile for global pandemic solutions. To isolate Nbs with potential neutralization breadth, recombinant RBD antigens from strains P.1 (isolated from Brazil) and B.1.617 (isolated from India) were used to screen a fully synthetic and highly diversified Nb phage display library^27^. After four rounds of reciprocal biopanning and phage ELISA, a panel of Nb binders was obtained. Total 18 Nbs were expressed in *Escherichia coli* and purified with one-step nickel affinity chromatography (Fig. 1a). The sequences of Nb complementary determining regions are listed in Table S1. To evaluate the neutralization breadth of these discovered Nbs, Spike-pseudotyped particle infection assay from four SARS-CoV-2 variants (B.1.1.7, B.1.341, P.1 and B.1.617) was performed. Encouragingly, several Nbs (Nb1, Nb2 and Nb15) demonstrated cross-protective activity at 0.33 μM, and each of them acted with a unique neutralization spectrum similarly or complementally (Fig. 1b). Thermal stability analysis showed that the Tm values range from 59.1 to 82.3 °C, with most of them above 70 °C (Fig. 1b).

**Fig. 1.**
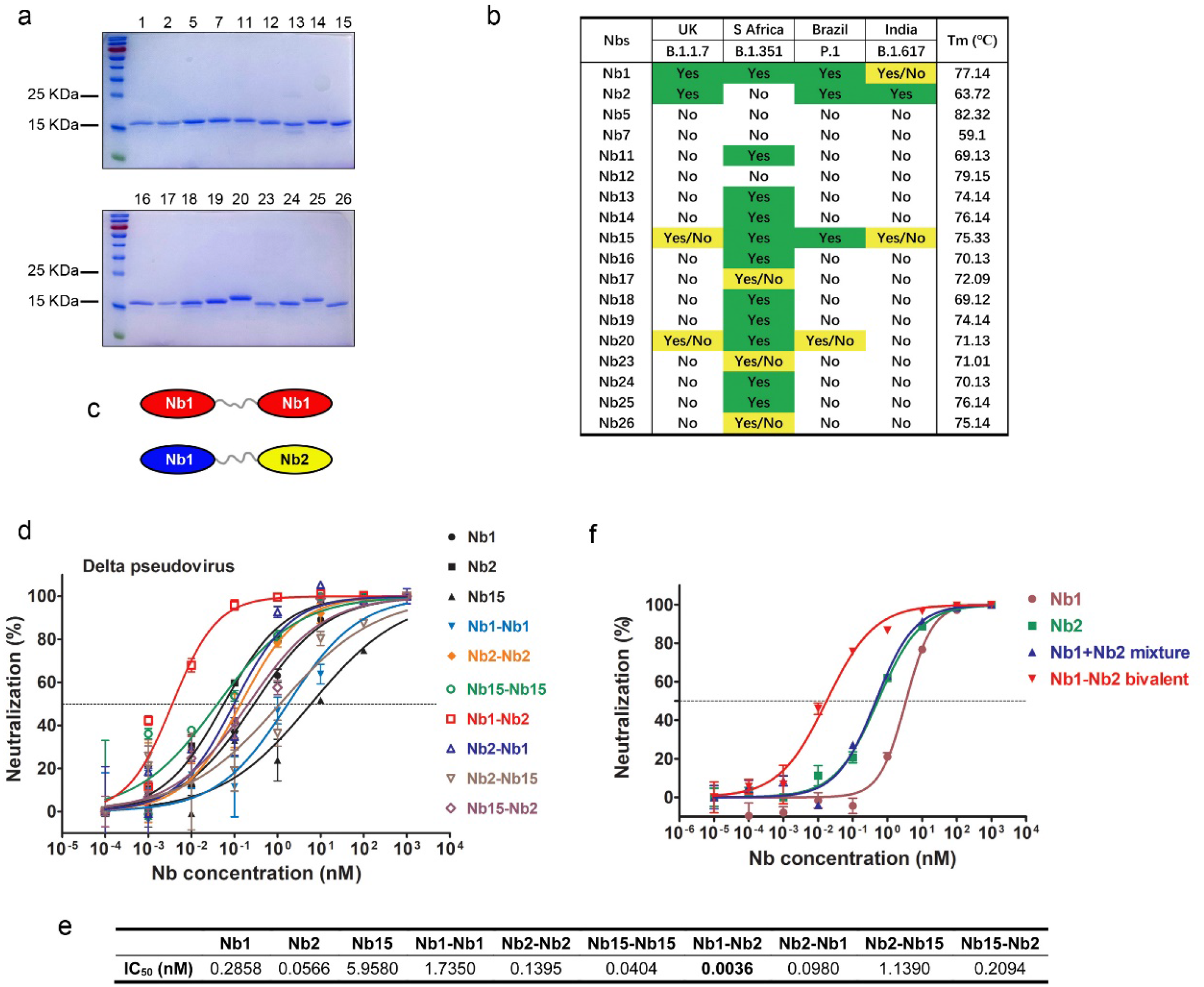
Screen and design of broad-spectrum neutralizing Nbs against SARS-CoV-2. **a** The purified recombinant proteins of SARS-CoV-2 RBD binding Nbs were separated by SDS-PAGE and stained with Coomassie Blue. **b** Nbs were incubated with the indicated SARS-CoV-2 variant pseudoviruses at a final concentration of 5 μg/mL (0.33uM) and inoculated into Huh7 cells. At 48 h post infection, luciferase activities were measured, and percent neutralization was calculated. Neutralization efficiency more than 90% was specified as Yes, 50%-90% as Yes/No, and less than 50% as No. Thermal stability of the purified Nbs were measured using circular dichroism spectra. **c** Schematic diagram for construction of homo-or heterodimeric Nbs. **d** Neutralization of SARS-CoV-2 Delta variant Spike-derived pseudovirus by various bivalent Nbs. The experiments were performed independently at least twice and similar results were obtained. One representative experiment was shown, and data were average values of three replicates (n = 3). **e** Summary of the half-maximal inhibitory concentration (IC_50_) values of Fig. 1D. **f** Pseudovirus neutralization activity of different Nb formulation.

Nb multimerization strategy can dramatically enhance the affinity and neutralization potency^28^. Thus, we designed a panel of homo- and hetero-dimeric Nbs by C-to N-terminus fusion expression with a flexible (GGGGS)_5_ linker sequence. Three Nbs (Nb1, Nb2 and Nb15) with relatively broad neutralization spectrum were chose as monomeric building blocks (Fig. 1c). Most bivalent Nbs showed improved neutralization activity against Delta variant-derived pseudovirus (Fig. 1d). Inspiringly, we found up to 15-79-fold activity increase for the heterodimer Nb1-Nb2 (IC_50_ =0.0036 nM) compared with the respective monomers (Fig. 1d & 1e). This enhanced neutralization potency could not be achieved by simply “cocktail” mixture formula of two monomers (Fig. 1f), suggesting a unique avidity binding mechanism to the trimeric spike.

We further determined the equilibrium-binding affinity (KD) of the monovalent and bivalent Nbs by BLI using multiple VOC derived RBD recombinant proteins, including the wild type of Wuhan isolate, Alpha (B.1.1.7, N501Y), Beta (B.1.351, K417N, E484K, N501Y), Gamma (P.1, L18F, K417T, E484K), Delta (B.1.617.2, L452R, T478K) and Kappa (B.1.617.1, L452R, E484Q) variants (Fig. 2). Nb1 bound to all VOC RBDs with a wide KD ranging from 4.4 to <0.001 nM. However, Nb2 showed selective affinity to wild type, Alpha, Delta, and Kappa RBDs with KD from 7.8-0.37 nM, but escaped from binding with Beta and Gamma RBDs. Through fusion connection, the bivalent Nb1-Nb2 demonstrated high affinity (less than or near 0.001 nM) to all RBDs (Fig. 2). These findings reveal that the bivalent format of Nb1 and Nb2 enhances the strength and breadth of its affinity to RBDs.

**Fig. 2.**
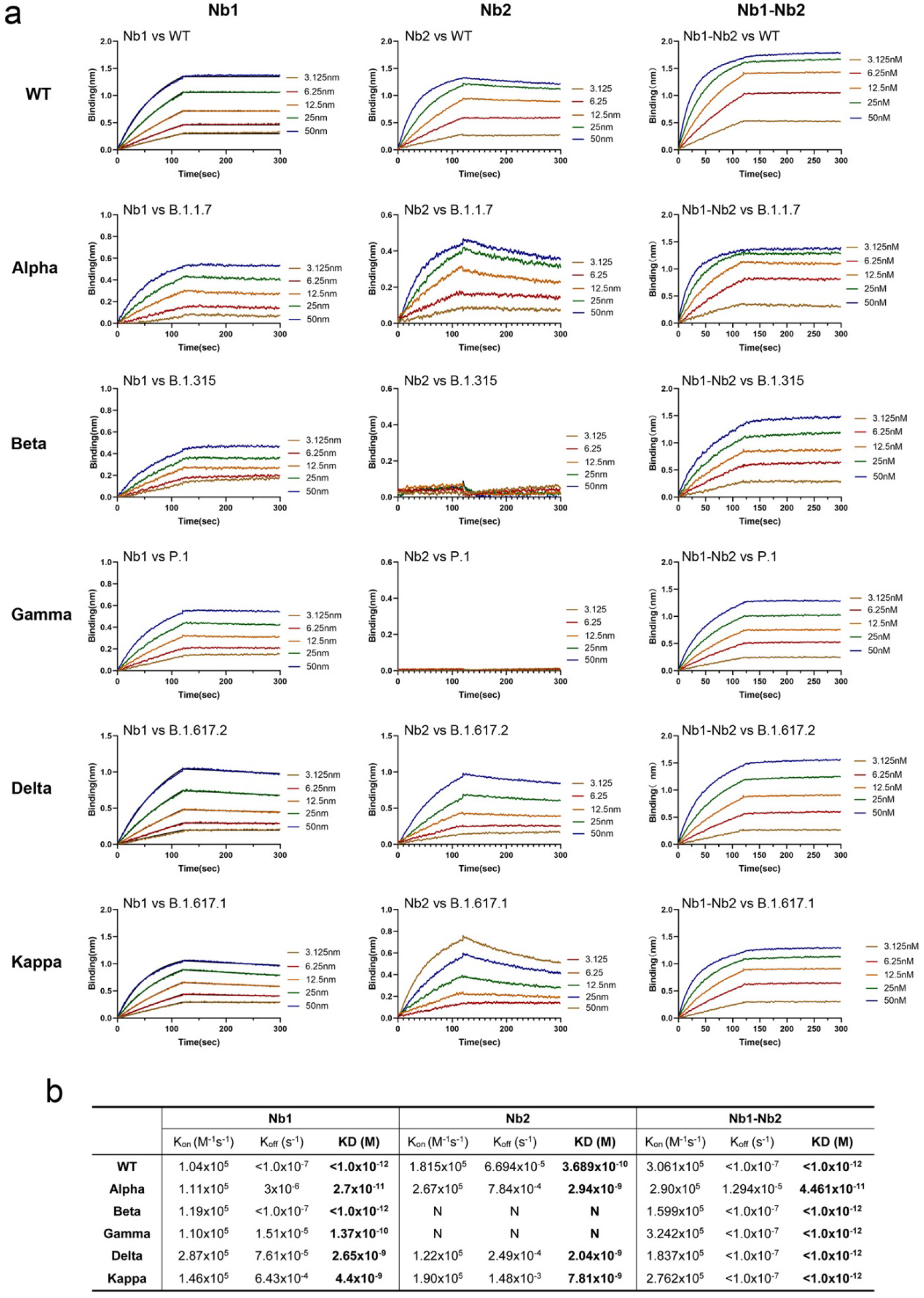
Binding affinity of Nbs against the RBDs from multiple circulating SARS-CoV-2 variants. **a** Fitted line plot showing the binding kinetic of Nbs with the immobilized receptor binding domain (RBD) proteins, measured using bio-layer interferometry (BLI). Recombinant RBD proteins were derived from SARS-CoV-2 WT, Alpha, Beta, Gamma, Delta or Kappa strain. The concentrations of Nb are shown in different colors. **b** Summary of BLI kinetic and affinity measurements. The equilibrium dissociation constant (KD), the association constant (K_on_) and the dissociation constant (K_off_) are presented. The assays without binding are marked as “N”.

### Epitope mapping using naturally occurring Spike mutants

SARS-CoV-2 will continue to evolve. SARS-CoV-2 Spike mutations, particularly in the RBD region, are strongly associated with the escape of antibody-mediated neutralization^29^. Currently, near a hundred mutation sites were found in Spike from the circulating SARS-CoV-2 isolates database GISAID^30^. The mutations in RBD, particularly in RBM, played critical roles for the increased transmission capability and neutralizing antibody resistance. To intensively analyze the effect of these mutations on the Nb neutralization potency, spike genes containing point mutations or deletions were generated and SARS-CoV-2 pseudoviruses were packaged. All the mutated pseudoviruses had a basic D614G substitution and the relative neutralization fold change to D614G was analyzed (Fig. 3a). For the Nb1 monomer, varying degrees of resistance were observed in a large proportion of mutant pseudoviruses, in which complete loss of neutralization was documented in the single point mutations A348S, N354D and T393P. Decreases more than 100-fold also included E471Q, E484K/Q, L452R/P681R and P681R/L452R/E484Q (Fig. 3a). A better situation happened with Nb2 monomer. The increased and decreased neutralization activities within 10-fold were evenly distributed among the tested mutations, though Nb2 suffered a complete loss of neutralization activity against N439K, E484K/Q as well as K417N (Fig. 3a). In general, the effect of mutations outside the RBD on the neutralization activity of the monomeric Nbs is less than that inside the RBD region. However, in sharp contrast, the bivalent Nb1-Nb2 exhibited an incredible neutralization spectrum width and enhanced potency. Among 64 mutated constructs, neutralization potency of Nb1-Nb2 was enhanced (2 to > 10-fold) in 46 of them, and a slight reduction (< 10-fold) was observed only in 15 mutants (Fig. 3a). The IC_50_ against various mutant pseudoviruses was summarized in Fig. 3b. These data imply that neutralizing antibodies with escape resistance can be designed by fusing two or more diverse Nbs.

**Fig. 3.**
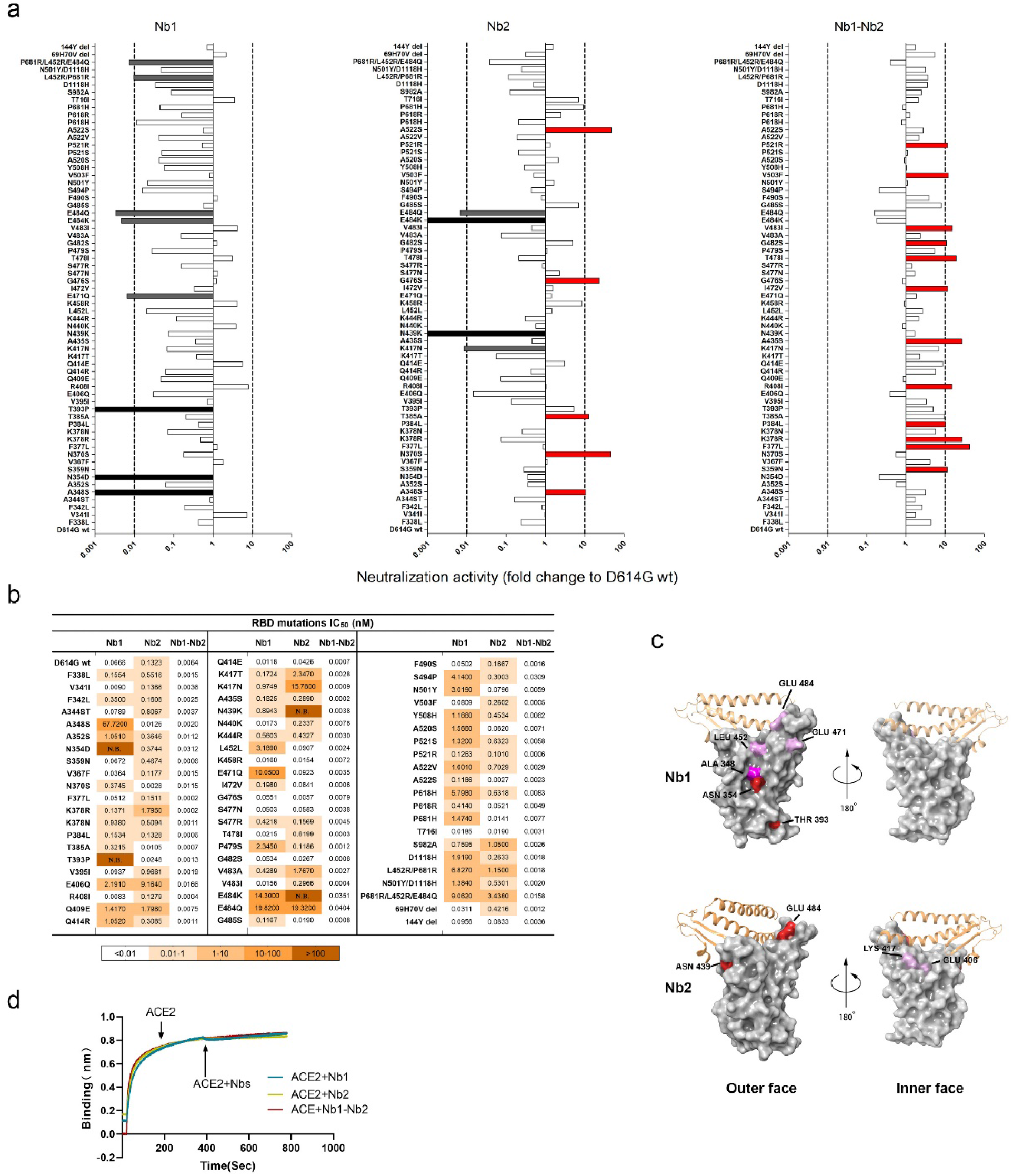
Epitope mapping using naturally occurring Spike mutants. **a** The SARS-CoV-2 pseudoviruses were packaged using more than 60 Spike variants identified from circulating viral sequences. The majority of mutations occur on RBD, including single amino acid substitution, combinational mutation and deletion. Neutralization activity conferred by Nb1, Nb2, and bivalent Nb1-Nb2 was evaluated. The x axis shows the ratio of IC_50_ of D614G pseudovirus/IC_50_ of indicated pseudovirus variant. When the ratio is greater than 1, the neutralization activity is increased, otherwise, the activity is decreased. The y axis shows the names of mutations. Data are represented as mean. All experiments were repeated at least twice. **b** IC_50_ values of indicated Nbs against SARS-CoV-2 mutation pseudovirus were calculated from data in Fig. 3a. **c** Location of critical amino acids on the RBD (PDB ID: 6M0J) region for Nb1 and Nb2. The key hot spots targeted by Nbs are shown in a color-coding pattern with resistant strength decending from red to pink. Both sides of RBD are shown from different angles. **d** Competition between Nbs and ACE2 for binding to the SARS-CoV-2 RBD. Octet sensors immobilized with the SARS-CoV-2 RBD were first saturated with ACE2 protein, and then exposed to the Nb1, Nb2 or Nb1-Nb2. The experiments were independently performed twice, and similar results were obtained.

Based on the above mutation analysis, we predicted the possible RBD epitopes for Nb1 and Nb2 by mapping the resistant hot spots on the surface of SARS-CoV-2 RBD (Fig. 3c). Currently, a consortium has been formed to define seven RBD communities (RBD-1 through RBD-7) that are bound by discovered neutralizing antibodies worldwide^18^. The antibodies in RBD-1 to RBD-3 target the top surface, namely RBM, and compete with ACE2. In comparison, antibodies in communities RBD-4/5 and RBD-6/7 bind to the outer and inner face of the RBD, respectively. Selecting antibodies for therapeutic cocktails benefits from this classification criteria. Interestingly, our prediction suggests that Nb1 recognizes an atypical RBD-4/5 mode with amino acids 348A/354N/393T as significant landmarks and 452E/471E/484E as potential influence sites (Fig. 3c). Nb2 adopts an approximate RBD-1/2/3 feature with amino acids 439N/484E/406E/417K as critical interaction points (Fig. 3c). The predicted binding sites of the two Nbs are both overlapping and separated, suggesting the RBD binding area could be enlarged through bivalent fusion of Nb1 and Nb2. To determine the neutralization mechanism, recombinant SARS-CoV-2 RBD was first immobilized on an AR2G biosensor and then saturated with ACE2. The addition of Nb1 or Nb2 to ACE2-saturated probe showed no complementary binding (Fig. 3d), which indicates that Nb1 or Nb2 have direct competition with ACE2 for binding to the SARS-CoV-2 RBD.

### Neutralization activity against multiple SARS-CoV-2 variants

As the SARS-CoV-2 continues to adapt and evolve in the human population, the dominant variants are also changing. To explore and compare the efficacy of the Nbs for neutralization of SARS-CoV-2 VOC, we first performed lentivirus-based pseudovirus infection assays. Seven pseudoviruses was produced to represent four VOC (Alpha, B.1.1.7; Beta, B.1.351; Gamma, P.1 and Delta, B.1.617.2) and three VOI (Lambda, C.37; Kappa, B.617.1 and Mu, B.621;). We found that bivalent Nb1-Nb2 broadly neutralized all pseudoviral variants with low IC_50_, ranging from 0.003 to 0.0865 nM (Fig. 4a & Fig. S1). However, the monomeric Nbs showed much lower activities, even loss of activity for Nb1 against Delta and Nb2 against Beta (Fig. 4a). These neutralization results were basically consistent with the affinity data (Fig. 2). Generally, monomeric Nbs provide a basic affinity and keep low activity. By designing a flexible bivalent strategy, the biparatopic Nb1-Nb2 can target two independent RBD epitopes and prevent or minimize viral escape.

**Fig. 4.**
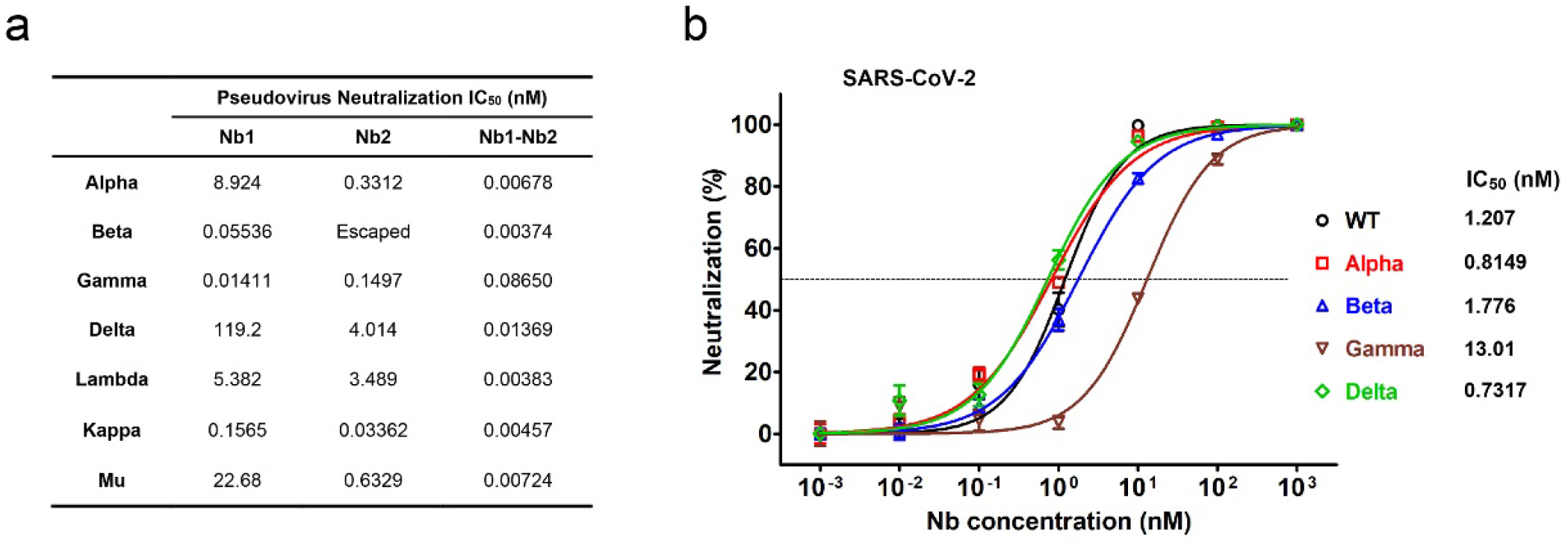
Neutralization of SARS-CoV-2 VOC and VOI by monomeric and bivalent Nbs. **a** Neutralization of pseudotyped SARS-CoV-2 variants by Nb1, Nb2 or Nb1-Nb2, respectively. Pseudovirus was pre-incubated with 10-fold serially diluted Nbs before inoculation of Huh7 cells. At 48 h post infection, luciferase activities were measured, and percent neutralization was calculated. The experiments were performed independently at least twice, and similar results were obtained. One representative data of one experiment were shown and data were average values of three replicates (n = 3). **b** Determination of neutralization efficacy of bivalent Nb1-Nb2 against recombinant SARS-CoV-2 GFP/ΔN trVLP. The infected cells were subjected to flow cytometry analysis for quantify the GFP fluorescence at 2 days post-infection. Error bars represent the standard deviations from three independent experiments (n = 3).

Neutralization assay of live SARS-CoV-2 (SARS-CoV-2 GFP/ΔN trVLP) that was constructed by reverse genetics was also performed (Fig. 4b)^31^. Bivalent Nb1-Nb2 neutralized wild type (WT) Wuhan strain SARS-CoV-2 GFP/ΔN trVLP with IC_50_ of 1.207 nM (0.036 μg/mL), and it had comparable activities to neutralize Alpha, Beta, Gamma and Delta live virus variants, with the IC_50_ around 0.8149 nM (0.024 μg/mL), 1.776 nM (0.054 μg/mL), 13.01 nM (0.390 μg/mL) and 0.7317 nM (0.022 μg/mL). Although the measured IC_50_ concentration in SARS-CoV-2 GFP/ΔN trVLP is higher than that in pseudovirus system, which may be due to sensitivity differences between the two virological tools, the trend of its broad-spectrum neutralizing activity is consistent. Importantly, the bivalent Nb1-Nb2 was effective against Beta (B.1.351) and Gamma (P.1) viruses, two of the most resistant variants leading almost complete loss of neutralization activity of the first generation RBM-associated antibodies^32^.

### Ultrapotent neutralization activity of the Fc-fused tetravalent biparatopic Nb

Although our bivalent Nb retains broad and relatively strong neutralizing activity, we hope to optimize its performance through further design. We constructed a human heavy chain antibody by fusing the human IgG1 Fc region to the C-terminus of bivalent Nb1-Nb2, making a tetravalent antibody through the disulfide bond formation in Fc hinge area (Fig. 5a). This optimized design can enhance antiviral activity, improve *in vivo* half-life and protein druggability. The tetravalent Nb1-Nb2-Fc was produced in Expi293F cells with supernatant yield > 20 μg per milliliter in a shaking flask (Fig. 5b). Most importantly, the tetravalent Nb1-Nb2-Fc exhibited extremely high neutralization potency against a panel of SARS-CoV-2 GFP/ΔN trVLP variants of concern. The neutralization IC_50_ values of the tetravalent Nb1-Nb2-Fc range from 0.0097 nM (0.0012 μg/mL) to 0.0987 nM (0.0118 μg/mL) depending on different variants, 32-183 folds increase to the corresponding bivalent Nb1-Nb2 (Fig. 5c-d). VOC and VOI derived pseudoviruses neutralization assays resulted in a similar activity enhancement (Fig. S2). In addition, the thermal stability was also satisfied for the Nb1-Nb2-Fc (Fig. S3).

**Fig. 5.**
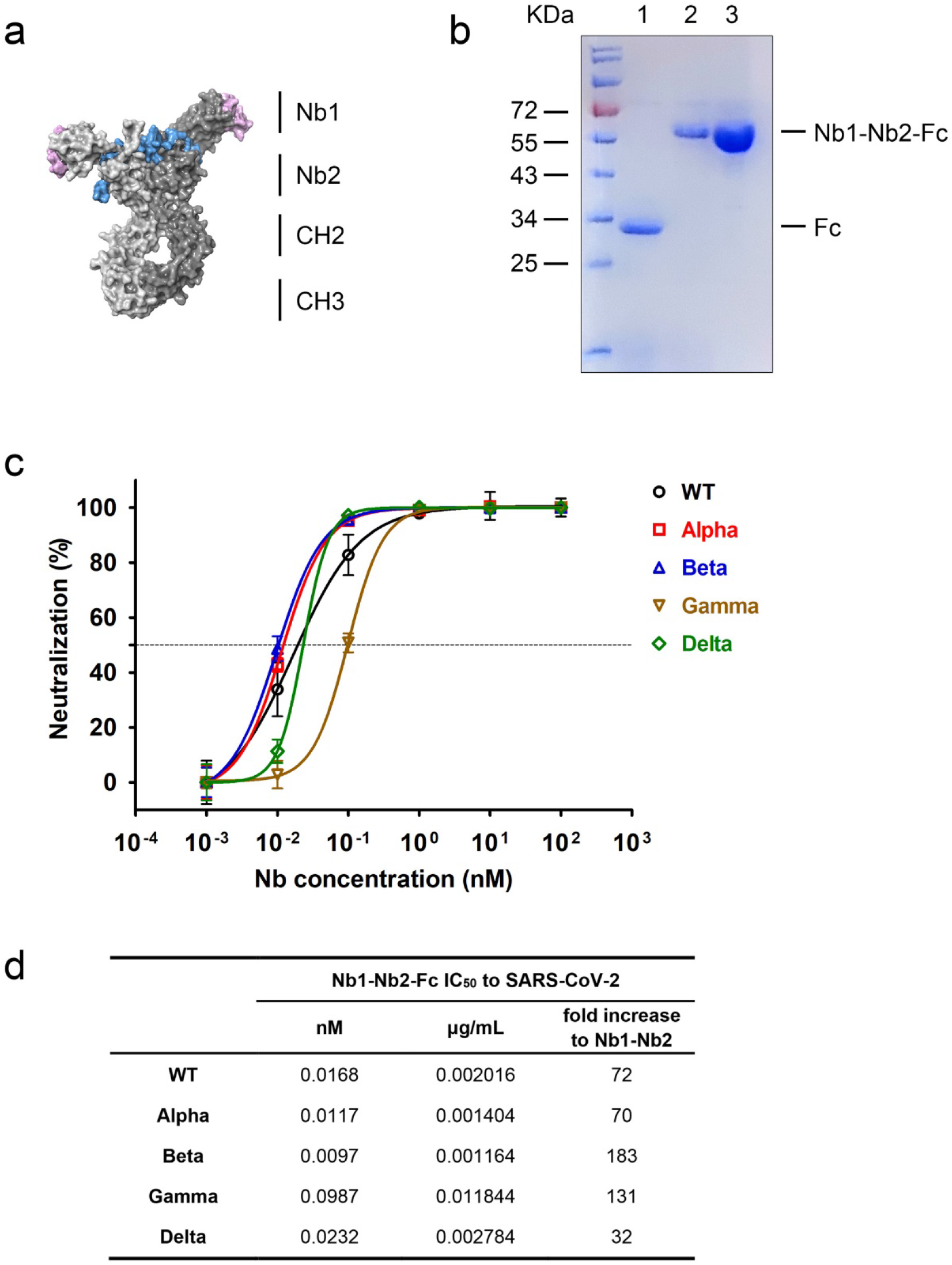
Enhanced neutralization potency by Fc-fused biparatopic Nb. **a** Schematic representation of the construction of Nb1-Nb2-Fc. Homology modeling of Nb1-Nb2-Fc was performed with SWISS-MODEL server. The structure is depicted as surface mode. The CDR regions were colored as pink for Nb1 and blue for Nb2. **b** Coomassie Blue staining of the Fc vector (lane 1) and Nb1-Nb2-Fc plasmid transfected Expi293F supernatants. Lane 3 shows the affinity purified Nb1-Nb2-Fc. Fc fusion to Nb1-Nb2 generates a heavy chain antibody with an approximate molecular weight of 60 kDa in reduced condition. **c** Neutralization of multiple SARS-CoV-2 GFP/ΔN trVLP variants with Nb1-Nb2-Fc. **d** Summary of neutralization IC_50_ value of Fc-fused Nb that was obtained in Fig. 5c. IC_50_ fold increases versus the corresponding non-Fc-fused bivalent Nb were calculated.

### Biparatopic Nb1-Nb2-Fc maintains high activity against the Omicron variant

The development of neutralizing antibody drugs for highly variable viruses has always been a challenge in the academy and industry. On 26 November 2021, WHO designated the variant B.1.1.529 a variant of concern, named Omicron. The Omicron RBD carries 15 mutations, most of which localize within the RBM region (Fig. 6a), resulting in reduced vaccine effectiveness and activity loss of many neutralizing antibodies^33,34^. We first measured the binding kinetic of different Nbs against Omicron RBD by BLI. Results showed that the tetravalent Nb1-Nb2-Fc demonstrated the strongest affinity with KD < 1.0×10^−12^ M, and the monomeric Nb1 and bivalent Nb1-Nb2 still maintained an ideal binding, though Omicron RBD completely escaped from binding with the monomeric Nb2 (Fig. 6b & 6d). Furthermore, to evaluate the neutralization potency of our Nbs against the Omicron variant, we packaged the pseudovirus harboring the Omicron Spike glycoprotein. It’s very encouraging that the tetravalent biparatopic Nb1-Nb2-Fc maintained potent neutralization activity against Omicron with IC_50_ around 0.0017 nM, which is comparable to other VOC and VOI pseudoviruses (Fig. 6d & Fig. S2). For the Omicron variant, there was no evidence showing the activity reduction for our biparatopic Nbs, either in terms of affinity or neutralization potency. All these data suggests that ultrapotent SARS-CoV-2 neutralization antibodies with mutation resistance can be obtained through optimized screening and reasonable design.

**Fig. 6.**
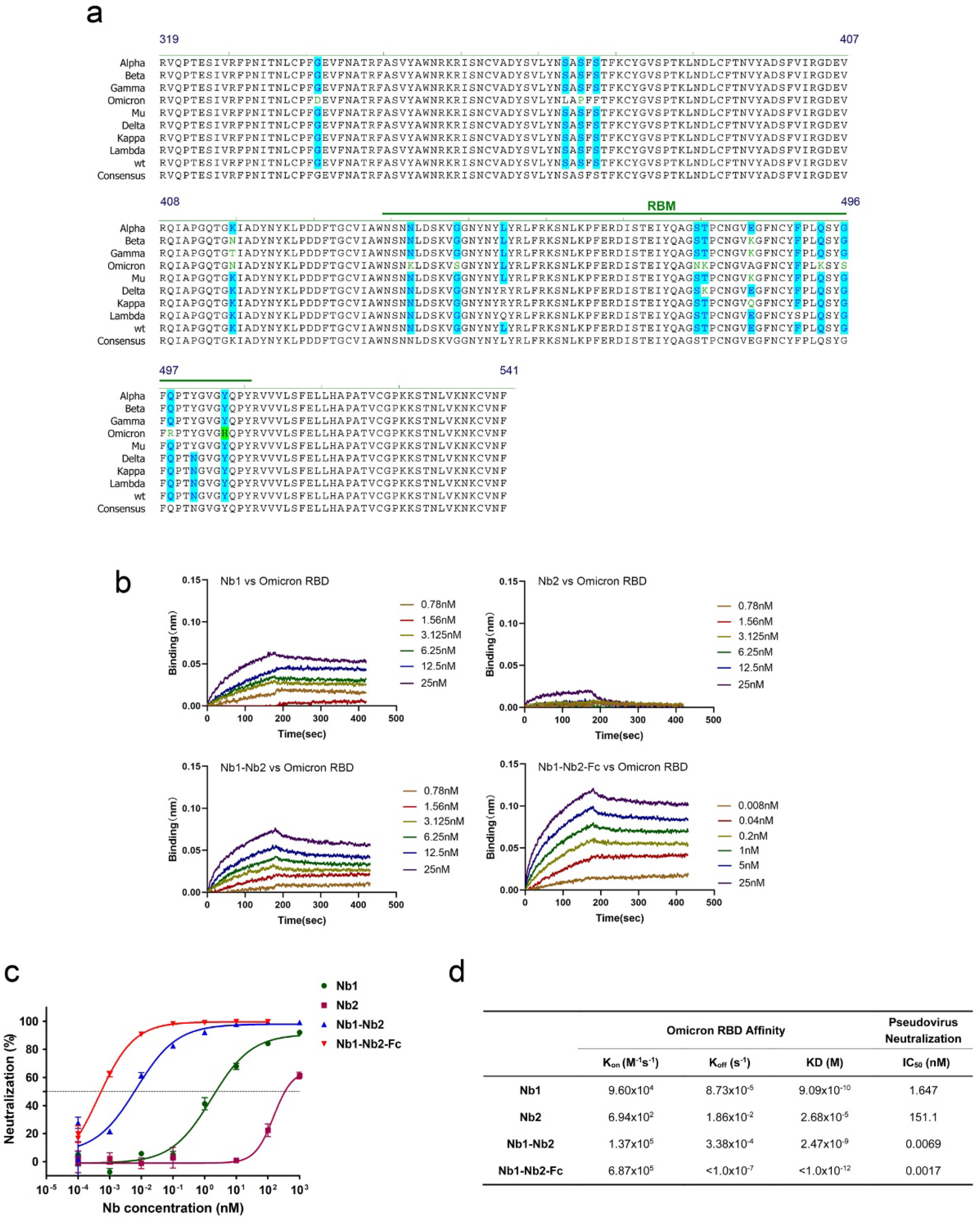
Neutralization of Omicron variant (B.1.1.529). **a** RBD amino acid sequence alignment from multiple VOC and VOI. Mutated points were colored. Consensus sequence was derived from the Wuhan isolate (wt). **b** Affinity analysis of four different Nbs against Omicron RBD with BLI. Fitted line plot showing the binding kinetic of four Nbs with the immobilized Omicron RBD. **c** Neutralization curve of Omicron pseudovirus by four Nbs. **d** Summary of binding kinetic and neutralization activity of four Nbs against Omicron variant. The K_on_, K_off_, KD and neutralization IC_50_ value are listed.

## Discussion

The COVID-19 vaccination rate has been accelerating globally, but SARS-CoV-2 transmission has no sign of stopping and the virus will continue to evolve. Recently, the variants Delta and Omicron have become the intensively concerned strains due to their unparalleled transmissibility. Several therapeutic antibody cocktails have been approved for postexposure treatment to reduce severe illness^12,13^. However, the potency of some antibodies is compromised by the emerging SARS-CoV-2 high-frequency mutation variants^32,35^, highlighting an urgent demand on developing next-generation antibodies with breadth and potency.

In this study, we identified a panel of SARS-CoV-2 neutralizing Nbs and designed a biparatopic heavy chain antibody Nb1-Nb2-Fc that targets the overlapping but distinct RBD antigenic regions and keeps cross-affinity with all of the SARS-CoV-2 RBD variants of concern that we tested. Nb1-Nb2-Fc broadly neutralizes pseudotyped viruses containing Spikes from the WHO designated variants Alpha, Beta, Gamma, Delta, Lambda, Kappa, Mu and Omicron, and other more than 60 representative circulating point mutated SARS-CoV-2 pseudoviruses. Neutralization against live SARS-CoV-2 variants was also confirmed and consistent with the pseudovirus model. As the virus is adapting to human and animal hosts and evolving rapidly, the antibody coverage and unique neutralization mechanism become critical to prevent and treat infection by emerging variants and to minimize the risk of viral escape. Our results indicate that the biparatopic heavy chain antibody is a promising one against the broad spectrum of variants currently being concerned.

Although more and more SARS-CoV-2 neutralizing antibody binding sites have been reported^36,37^, including NTD, the most effective epitopes localize on RBD due to its natural role to bind human ACE2. SARS-CoV-2 RBD has been structurally defined into seven “core” antibody-binding communities. Generally, antibodies from communities RBD-1 through RBD-4 are relatively more potent, but highly susceptible to neutralization escape by mutations. Whereas RBD-5 through RBD-7 binding antibodies often have lower potency but are more resistant to escape. Although we attempted to use crystallography and cryo-electron microscopy to elucidate the details of the interaction between the identified Nbs and RBD, unfortunately, valuable information was not obtained. Using highly extensive RBD mutagenesis and pseudovirus techniques, we identified the key amino acids resistant to Nb neutralization. Based on the mapped epitope sequences, Nb1 and Nb2 demonstrated different binding sites but overlapped to some extent. Nb1 recognized sites localizes on the outer face of the RBD, but Nb2 binds mainly to the RBM. Interestingly, Glutamic Acid at position 484 is the shared landmark amino acid for both Nb1 and Nb2. For the monomeric Nb1 and Nb2, their neutralization potency was significantly lowered or even lost when several key amino acids were altered. However, benefiting from the rational design of biparatopic strategy, our tandem fusion form of Nb1 and Nb2 can efficiently neutralize all mutant pseudoviruses and variants of concern, some of which are completely resistant to the monomeric Nbs.

In recent years with the SARS-CoV-2 pandemic, many attentions have been turned to Nbs, also known as single-domain antibody or VHH, which are derived from camelids and easier to produce. SARS-CoV-2 neutralizing Nbs have been discovered and reported by several laboratories, including ours^27,38^. Because of their small size and flexible combination for multimers, Nbs are becoming powerful weapons against pathogens. Multivalent Nbs have been documented for several viruses with much stronger neutralization potency than single Nbs^25^, and multivalent antibodies that bind two epitopes also prevent the emergence of viral escape mutants^39^. It’s worth pointing out that the neutralization activity is increased for tens to 2 log fold in the tetravalent Nb1-Nb2-Fc context, which functions as a heavy chain antibody with a double punch against SARS-CoV-2 in each arm through biepitopic binding. The tetravalent Nb1-Nb2-Fc is evident with high yield in mammalian cells and durable stability at 37°C, potentiating the application for either an injectable formula or administration by inhalation.

Taken together, the results presented here for Nb-based neutralizing antibody development, offers a detailed pipeline and strategy to combat with emerging SARS-CoV-2 variants with super wide neutralization breadth. Moreover, these findings indicate that Nb1-Nb2-Fc is a promising candidate for clinical development and could be stockpiled as part of a pandemic readiness toolbox.

## Materials and Methods

### Cells and reagents

The HEK293T (human kidney epithelial) cells were obtained from China Infrastructure of Cell Line Resource (Beijing, China). The human hepatoma cell line Huh7 was obtained from Apath, Inc (Brooklyn, NY, USA) with permission from Dr. Charles Rice (Rockefeller University). The Expi293F cells were purchased from ThermoFisher (Waltham, MA, USA). The cells were maintained in Dulbecco’s modified Eagle’s medium (ThermoFisher) supplemented with 2-10% fetal bovine serum (FBS, ThermoFisher), non-essential amino acid, penicillin and streptomycin. Recombinant RBD and ACE2 proteins were purchased from Sino Biological (Beijing, China). HRP/anti-CM13 monoclonal conjugate was from GE Healthcare (Boston, MA, USA).

### Screen of nanobody library

A synthetic nanobody phage display library with high-diversity was prepared as previously described^27^. Screening for nanobodies was performed by panning in both immunotubes and with magnetic bead-conjugated antigen, using SARS-CoV-2 variants P.1 and B.1.617 derived recombinant RBD proteins. Briefly, for the 2nd and 4th panning rounds, the purified SARS-CoV-2 RBD proteins were coated on Nunc MaxiSorp immuno tubes (ThermoFisher) at 5μg/mL in PBS overnight. For the 1st and 3rd panning rounds, RBD protein was first biotinylated with EZ-Link™ Sulfo-NHS-LC-Biotin (ThermoFisher) and then selected with streptavidin-coated magnetic Dynabeads™ M-280 (ThermoFisher). The panning was performed according to a standard protocol^27^. After 4 rounds of panning, phage ELISA identification was performed with 960 individual colonies using Anti-CM13 antibody in the plates coated with recombinant RBDs. The absorbance was measured using a SpectraMax M5 plate reader from Molecular Devices (San Jose, CA, USA). The positive clones were sent for sequencing. After sequence alignments, the distinct sequences were chosen for protein expression.

### Expression and purification of nanobodies

Full-length sequences of selected nanobodies were PCR amplified and cloned into the NcoI/XhoI sites of the pET28b (Novagen, Sacramento, CA, USA) and transformed into BL21(DE3) chemically competent E. coli. The expression of recombinant nanobodies was induced by adding IPTG to a final concentration of 0.3 mM after culture had reached OD_600_=0.5-0.6 and grown over night at 25°C. The nanobodies were fused with a His-tag at C-terminus and purified over Ni Sepharose 6 Fast Flow (GE Healthcare) and eluted with 400 mM imidazole. Affinity purified sdAbs were dialyzed against PBS to eliminate imidazole.

### Construction of bivalent and Fc-fused nanobodies

To improve the neutralization activity of Nbs, we constructed dimeric nanobodies with various combinations and a (GGGGS)_5_ linker was introduced between the two monomers. The recombinant bivalent nanobodies were produced in E *coli*. and a His-tag was designed to facilitate purification. In addition, the sequence of dimeric Nb1-Nb2 was cloned into a mammalian expression vector under the control of hEF1-HTLV promotor and fused with N-terminal interleukin-2 signal peptide and C-terminal Fc region, comprising the CH2 and CH3 domains of human IgG1 heavy chain and the hinge region. Maxiprepped plasmids were transiently transfected into Expi293F cells (Thermofisher) and the cells were further cultured in suspension for 2-3 days before harvesting antibody-containing supernatant. Fc-fused nanobody was prepared with prepacked HiTrap® Protein A HP column (GE Healthcare). The produced Fc-fusion protein was analyzed by SDS-PAGE using standard protocols for dimerization, yield and purity measurement.

### Pseudotyped virus and neutralization assay

To produce SARS-CoV-2 pseudovirus, HEK293T cells were seeded 1 day before transfection at 2.5×10^6^ cells in a 10-cm plate. The next day, cells were transfected using Lipofectamine 2000 (ThermoFisher). The plasmid DNA transfection mixture (1 ml) was composed of 15 μg of pNL-4.3-Luc-E-R- and 15 μg of pcDNA-SARS-CoV-2-S that was purchased from Sino Biologicals and reconstructed by deletion of 18 amino acid cytoplasmic tail. A nonenveloped lentivirus particle (Bald virus) was also generated as negative control. Sixteen hours after transfection, the media was replaced with fresh media supplemented with 2% FBS. Supernatants containing pseudovirus were typically harvested at 36–48 h after transfection and then filtered through a syringe filter (0.22μm) to remove any cell debris. The pseudovirus was freshly used or allocated and frozen at -80°C. To conduct the virus entry assay, 1×10^4^ Huh7 cells were seeded in each well of a 96-well plate at 1 day prior to transduction. The next day, 100 μL of supernatant containing pseudovirus was added into each well in the absence or presence of serially diluted Nbs or human IgG1 Fc-fused Nb. Forty-eight hours after transduction, the cells were lysed in 100 μL of passive lysis buffer and 50 μL lysate was incubated with 100 μL of luciferase assay substrate according to the manufacturer’s instructions (Promega, Madison, WI, USA).

Substitutions of the residues at the sites selected for mutagenesis were based on the pcDNA3.1-SARS-CoV-2-S (GenBank: MN_908947), which was purchased from Sino Biologicals and reconstructed by deletion of 18 amino acid cytoplasmic tail. Following the procedure of circular PCR, 15 to 20 nucleotides before and after the target mutation site were selected as forward primers, while the reverse complementary sequences were selected as reverse primers. Site-directed mutagenesis was induced with a commercialized KOD-Plus mutagenesis kit (TOYOBO, Cat. No.SMK-101). The mutations were confirmed by DNA sequence analysis (Rui Biotech, Guangzhou, China). The primers for the specific mutation sites are in Table S2. For the variants derived pseudovirus, the spike genes were codon optimized, synthesized, and cloned into pCAGGS vector. The profile of amino acid changes compared to the wild-type virus (Genbank QHD43416.1) for each variant are listed in Table S3.

### Production of genetic complementation SARS-CoV-2 (SARS-CoV-2 GFP/ΔN trVLP)

A nucleocapsid (N)-based genetic complementation system for production of SARS-CoV-2 at BSL-2 laboratory was described previously^31^. Briefly, cDNAs (for multiple variants) of SARS-CoV-2 GFP/ΔN were synthesized. The N gene is replaced with the gene of green fluorescent protein (GFP). RNA transcripts were *in vitro* transcribed by the mMESSAGE mMACHINE T7 Transcription Kit (ThermoFisher Scientific) and transfected into Caco-2-N cells by electroporation. The produced SARS-CoV-2 can be amplified and titrated in Caco-2-N cells. Serially diluted antibodies were mixed with SARS-CoV-2 and inoculated into Caco-2-N cells. The infection efficiency was measured by flow cytometry analysis at 48 h post infection.

### Biolayer interferometry (BLI) measurement

Antibody affinity analysis was conducted by ForteBio Octet RRD 96 system. The VOC/VOI derived RBD recombinant proteins (Sino Biological, Cat: 40592-V08H/V08H82/V08H85/V08H86/V08H88/V08H90) were diluted in 10 mM Acetate pH 5.5 buffer at a density of 10μg/ml. The Amine Reactive 2^nd^ Generation biosensors surface was activated with EDC and NHS, then immobilized the RBD proteins for 5 min. Following 10 s of baseline in kinetic buffer (KB: 1× PBS, 0.01% BSA, and 0.02% Tween-20), the loaded biosensors were dipped into serially diluted (3.125–50 nM) nanobodies for 120 s to record association kinetics. The sensors were then dipped into kinetic buffer for 180 s to record dissociation kinetics. Kinetic buffer without antibody was set to correct the background. The Octet Data Acquisition 9.0 was used to collect affinity data. The mean Kon, Koff, and apparent KD values were calculated using an equation globally fitted to a 1:1 binding kinetic model and using the global fitting method.

### Epitope competition-binding Study

For the ACE2 competition assay, the RBD-immobilized biosensors were then dipped into the wells containing 100 nM of ACE2 for a 360-s association period. The sensors were then transferred to wells containing 100nM ACE2 or 100nM ACE2 +100nM Nb samples and incubated for 400s. For all BLI assays, data analysis was performed using Octet data analysis software version 11.0 (Pall FortéBio).

### Stability tests

Antibody samples diluted in PBS (1 mg/mL) were filtrated and sealed in a 1.5 mL Eppendorf tube and stored at 37°C for 3 or 6 days. At the end of the storage period, samples were centrifuged (10,000× g) for 10 min and neutralization activities were evaluated using pseudovirus.

### Circular Dichroism measurements

CD spectral data of the protein solution was obtained using the Spectra Measurement program on a Jasco J-815 CD spectrometer equipped with a 1.0 mm path length unit. HBS solution with 20 mmol/L concentration was mixed with Nano antibodies separately so that the final concentration was 15 μmol/L. The wavelength range from 200 nm to 250 nm was scanned and far-ultraviolet spectrum data was collected. SpectraManger software was used to process the collected data to obtain the content of the circular chromatogram of each system. Temperature regulation was carried out using the Variable Temperature Measurement program. A data pitch of 0.1 nm and bandwidth of 1 nm was used. Heat-induced unfolding was recorded at 208 or 218 nm, and a heating rate of 0.5°C/min was used.

### Antibody-escape sites visualization

The RBD and ACE2 binding crystal structure (PDB: 6M0J) is represented by a surface pattern. The antibody-escape amino acids on RBD are colored at each site. Different degrees of escape are indicated in different colors. Red represents complete escape, pink represents moderate escape, and light pink represents weak escape. Interactive visualizations of the escape maps and their projection onto the ACE2-bound were created using dms-view (https://dms-view.github.io/docs/).

### Statistics and reproducibility

Data were analyzed using GraphPad Prism 6.01 (GraphPad Software, San Diego, CA, USA). The values shown in the graphs are presented as means ± SD. One representative result from at least two independent experiments was shown. Antibody neutralization experiments usually use three to four duplicated wells for each treatment. The infectivity data were first inversed to neutralization activity. Each neutralization data set was normalized by the background control (no virus) to define the real value for 100% neutralization. After transformation to neutralization, the lowest concentration point of antibody treatment was set to 0% neutralization. Then, a 4-parameters neutralization nonlinear regression model was fitted to report IC_50_ values. All experiments were performed independently at least twice and similar results were obtained. One representative data of one experiment were shown.

## Data Availability

The source data underlying Figs. 1a, 1d, 1f, 4a, 4b, 5c, 6c are provided as a Source Data file. The sequences of Nb CDRs are listed in Table S1. All other data are available from the corresponding author upon reasonable requests.

## Competing interests

A patent application has been filed on the nanobodies reported in this study.

## ACKNOWLEDGEMENTS

This work was supported by CAMS Innovation Fund for Medical Sciences (2021-I2M-1-038) and National Natural Science Foundation of China (81871667 and 82002153). Drs Xiangxi Wang, Sheng Cui, Zhijian Li, Lei Wang and Han Wen provided help and advice on structural biology analysis.

